# Somato-Motor Network Neural Connectivity Correlates with Visuomotor Adaptation

**DOI:** 10.1101/2025.05.19.654905

**Authors:** Yanlong Song

## Abstract

Humans show tremendous individual differences in learning motor skills. Prediction of individual differences in motor learning is of theoretical significance and practical relevance. Brain as the central neural apparatus supporting motor control and learning has attracted considerable research efforts on examining brain functional and structural predictors of individual differences in motor learning. Many previous studies applied a resting state (RS) approach in recording brain activity, selected the contralateral primary motor cortex (M1) a priori as a seed or region of interest (ROI), and reported that functional connectivity between M1 and other brain regions predicted individual differences in tasks mostly on motor sequence learning. However, task-evoked brain neural activity and connectivity that represent different brain dynamics from spontaneous brain dynamics have not been studied in this line of research. This study presents findings on RS and finger-tapping evoked cortical somato-motor network (SMN) neural connectivity correlates with individual performances in a visuomotor adaptation (VMA) task, based on analysis of multimodal data from the Cam-CAN study. SMN nodes were initially localized through finger-tapping evoked brain magnetic fields as sampled with magnetoencephalography (MEG). It was found that SMN node specific neural connectivity strength at the contralateral primary somatosensory cortex (S1), supplementary motor area (SMA), and dorsal premotor cortex (PMd) during different stages around finger tapping but not in RS were significantly correlated with individual mean target errors in the final adaptation stage. The results imply critical roles of somatosensory and secondary motor areas in human motor learning.

## Introduction

Humans have marvelous capacities of performing skilled movements but also tremendous individual differences in learning motor skills. One person may need only days of practice to master a motor skill while another person may take months of practice to achieve a comparable level of motor performance. Prediction of individual differences in motor learning is of theoretical significance and practical impacts; for instance, theoretically, it may reveal determinants of motor abilities, and practically, it may optimize design and administration of practice schedules in sports training or motor rehabilitation.

Factors that are predictive of individual differences in motor learning can be categorized into during-practice factors and outside-practice factors according to Ranganathan et al. (2022), who reported that 10.8% of published studies that examined predictors of individual differences in motor learning tested during-practice factors and 89.2% of published studies examined outside-practice factors. Research on during-practice factors may uncover predictors of individual differences specific to a motor task while research on outside-practice factors may lead to discovery of predictors of individual differences across different motor tasks. The generalizability of a predictor is of importance since it implicates a general ability or trait that may support acquisition of diverse motor skills across contexts.

Brain, the central neural apparatus to learn and control skilled movements, has attracted considerable research efforts on searching for brain structural and functional correlates with individual performances in motor learning. Earlier studies focused on the contralateral primary motor cortex (M1), a core region that underpins motor control and motor learning (Lee et al., 2022). Several factors measuring M1 functions and structures, including neural oscillations in alpha and beta frequency bands (Krause et al., 2016; Pollok et al., 2014), M1 neural excitability as derived from transcranial magnetic stimulation (TMS) of M1 and TMS-evoked electromyography (EMG) responses (Hirano et al., 2015; McGregor et al., 2018), GABA concentration at M1 (Dupont-Hadwen et al., 2020; Kolasinski et al., 2019), movement-related gamma oscillation power increase at M1 (Zich et al., 2025), and gray matter volume and white matter microstructure (Sampaio-Baptista et al., 2014; Tomassini et al., 2013), are correlated with individual performance differences in diverse laboratory motor learning tasks.

As brain connectome gains momentum, research that examines brain connectivity predictors of individual differences in motor learning has been continuously growing. Electroencephalography (EEG), magnetoencephalography (MEG), and functional magnetic resonance imaging (fMRI) have been applied to measure brain connectivity in resting states (RS). Extending earlier studies with a focus on M1, many studies employed an approach of selecting M1 a priori as a seed or region of interest (ROI) to estimate RS connectivity between M1 and other brain regions and then correlate M1-seeded connectivity with individual motor learning performances. For instance, MEG beta-band (13-30Hz) RS neural connectivity between M1 and several regions including temporal gyrus, supramarginal gyrus, postcentral gyrus, precentral gyrus, and parietal lobule was associated with performances of subsequent motor sequence learning (Sugata et al., 2020); MEG 0.5 Hz to 45 Hz RS connectivity between M1 and putamen or cerebellum predicted motor sequence learning differences in younger adults while RS connectivity between M1 and dorsolateral prefrontal cortex, precuneus, or anterior cingulate cortex predicted learning differences in older adults (Mary et al., 2017); EEG alpha-band (8-12Hz) (Mottaz et al., 2024) and 0.1Hz to 50 Hz (Wu et al., 2014) RS connectivity between M1 and all other brain regions predicted individual learning of motor sequences or visuomotor tracking; fMRI RS functional connectivity between M1 and somatosensory cortex was associated with individual force-field motor adaptation gains after a period of observation (McGregor & Gribble, 2017); fMRI RS connectivity between M1 and cerebellum were correlated with individual motor sequence learning differences (Bonzano et al., 2015). Beyond M1, RS connectivity estimated from supplementary motor area (SMA) (Bonzano et al., 2015), somatosensory cortex (McGregor & Gribble, 2017), premotor cortex (PMC) (Hardwick et al., 2015), prefrontal cortex (Mizuguchi et al., 2019; Omurtag et al., 2025), striatum and mediotemporal lobe (Mottaz et al., 2024), cerebellum (Baldassarre et al., 2021), and superior parietal cortex (Manuel et al., 2018) were predictive of individual differences in diverse motor learning tasks.

Most previous studies that examined functional connectivity of M1 and other brain regions and their associations with individual differences in motor learning applied an RS procedure to record brain activities, had relatively small samples of participants (most less than 30), and employed an approach of selecting ROIs based on a standard anatomical atlas. In contrast to spontaneous RS brain activity, brain activity evoked by specific stimuli or motor responses are time-locked activity, which represent different brain dynamics and show higher inter-individual consistency than non-time-locked spontaneous activity (Arviv et al., 2015). Brain functional network analysis revealed a task-general network architecture highly similar to RS network architecture and task-specific network changes evoked by specific tasks (Cole et al., 2014). However, task-evoked brain neural activity and connectivity have been rarely studied in the line of research on prediction of individual differences in motor learning.

The present study was targeted to elucidate brain somato-motor network (SMN) neural connectivity correlates with individual differences in visuomotor adaptation (VMA), in which visual feedback of a moving virtual cursor that represents the reaching hand was rotated by some degree to assess adaptation of an old visuomotor association and formation of a new visuomotor association. SMN supports preparation and execution of voluntary movements and processing of somatosensory information. Overlap has been demonstrated between SMN derived from RS functional imaging and brain activation maps evoked by motor tasks (Beheshtian et al., 2021; Schneider et al., 2016). It was hypothesized that SMN global neural connectivity and node specific neural connectivity would be predictive of individual differences in VMA. Individual VMA performance data and brain neural responses recorded with MEG during RS and visual-auditory cued finger tapping (VACFT) from a public data repository Cam-Can were accessed and analyzed to test the hypothesis. MEG provides direct measures of whole brain neural activities with superior temporal resolution and sufficient spatial resolution (Lopes da Silva, 2013) and thus is an ideal technology to assess brain network neural connectivity. Key nodes in SMN were initially functionally localized from MEG recordings in the VACFT task. Then neural connectivity among the functionally localized key nodes in RS and time locked to VACFT were estimated and correlated with individual adaptation performances. It was found that SMN node specific neural connectivity strength during different stages in the VACFT but not in RS were significantly correlated with individual final adaptation performance.

## Methods

### Ethics statement

Data used in this study were sampled from a public data repository for the original Cambridge center for Ageing and Neuroscience (Cam-CAN) dataset (available at http://www.mrc-cbu.cam.ac.uk/datasets/camcan/), (Shafto et al., 2014; Taylor et al., 2017). The Cam-CAN project was approved by the University of Cambridge, Cambridge, UK. All participants provided written informed consent to participate in the Cam-CAN project.

### Data Description

The Cam-CAN dataset contains data from about 700 participants (100 per decade from 18 to 89), who participated in multi-modal tasks including MEG, structural and functional Magnetic Resonance Imaging (MRI), and multiple behavioral experiments. This study stratified participants who completed T1-weighted brain MRI scans, MEG recordings during RS and a VACFT task, VMA, and hand dominance assessment into three age groups including 18 to 39 years old (YO), 40 to 64 YO, and 65 to 89 YO, then randomly selected 32 right-handed participants in each age group (96 in total).

### Data Acquisition

Taylor et al. (2017) described details of data acquisition in the Cam-CAN dataset. In summary, T1-weighted brain MRI scans with a spatial resolution of 1mm x 1mm x 1mm were acquired using the three-dimension magnetization-prepared rapid gradient-echo (MPRAGE) sequence. MEG data were recorded with an Elekta whole-head SQUID sensor array with a sampling rate of 1000Hz in RS in which participants rested with eyes closed and a sensorimotor task in which participants responded to a visual-auditory cue with a single tapping of the dominant hand index. Simultaneous electrocardiogram (ECG) and electrooculography (EOG) were also recorded during MEG recordings. VMA was measured in a fast-pointing task of 192 trials with veridical, 30º rotated, and veridical again visual feedback of a hand-and-stylus-controlled computer cursor during shooting of a virtual target. The mean of the target errors across the final 25% of the 120 adaptation trials was calculated to measure the achieved adaptation level in a participant by the Cam-CAN group. Hand dominance was assessed with the Edinburgh Handedness Inventory.

### MEG Data Analysis

MEG recordings available for this study were initially filtered to reduce environmental noise and compensate for head movements with MaxFilter (MEGIN, Finland) using the temporal extension of signal-space separation by the Cam-CAN group. The MEG recordings sampled from the 96 participants were then processed with the open-access toolbox Brainstorm (version: Aug-2024) (Tadel et al., 2011). MEG data analysis steps included preprocessing, evoked-response analysis, source activation mapping, and neural connectivity analysis.

Preprocessing of MEG signals started from reviewing of continuous signals and removal of bad segments and bad channels manually, followed with direct current offset removal, 1 to 100 Hz band-pass filter, and 60Hz notch filter. MEG artifacts due to heartbeats and blinks were automatically detected from ECG and EOG recordings, then removed with signal-space projection (SSP) (Tesche et al., 1995) and independent component analysis (ICA) (Bell and Sejnowski, 1995) algorithms by calling corresponding Brainstorm default functions.

Preprocessed continuous sensorimotor MEG signals were segmented into event-locked trials with a duration of 3200 ms, 1600 ms prior to finger tapping onset and 1600 ms post finger tapping onset to assess sensor-level evoked responses time-locked to tapping of the dominant hand index in response to a visual-auditory cue. The average of clean trials, namely tapping-evoked fields, was computed for each participant. Similarly, preprocessed continuous RS MEG signals were segmented into trials with a duration of 3200 ms but without referencing to any event and averaging.

Individual cortical neural source activations underlying sensor tapping-evoked fields or RS fields were reconstructed via application of a noise-normalized source inversion solution dynamic Statistical Parametric Mapping (dSPM) (Dale et al., 2000). MEG sensors were co-registered to an individual participant’s 3D head model that was simulated with canonical surfaces (cortex, inner skull, outer skull, and scalp) that were generated from individual T1-weighted brain MRI scans. A boundary element model (BEM) was constructed from the co-registration with the OpenMEEG toolbox (Gramfort et al., 2010) called by Brainstorm to solve the forward problem, which is about computation of sensor magnetic fields for a putative source configuration. The dSPM source inversion of neural activations underlying tapping-evoked fields or RS fields was estimated for the cortex surface (~15,000 vertices) with both gradiometers and magnetometers included, dipole orientations constrained to the cortex, and noise covariance matrix estimated from the baseline interval between −1500 ms and −1000 ms prior to tapping for tapping-evoked fields or an empty room recording for RS fields.

Group-level significant cortical neural activations underlying tapping-evoked fields were analyzed in a common MNI152 template cortex source space (Fonov et al., 2009) by projecting an individual’s vertex-wise mean dSPM estimations across trials to the MNI152 template to lessen volume conduction and individual structural variation influences on MEG sensor-level data (Winter et al., 2007). Individual MNI-normalized cortical dSPM estimations were further averaged across participants to form the group-level mean vertex-wise dSPM estimations, which displayed five components before and after tapping (Fig. 1). Group-level significant cortical neural activations underlying tapping-evoked fields were estimated in the five components of the group-level mean vertex-wise dSPM estimations by paired-sample permutation t tests to compare differences between individual’s mean of MNI-normalized dSPM estimations within each component to mean of MNI-normalized dSPM estimations within the pre-tapping baseline. Group-level cortical activations estimated with dSPM were deemed as significant if p < 0.001 from a t test at any component (uncorrected for multiple comparisons).

**Fig 1.**
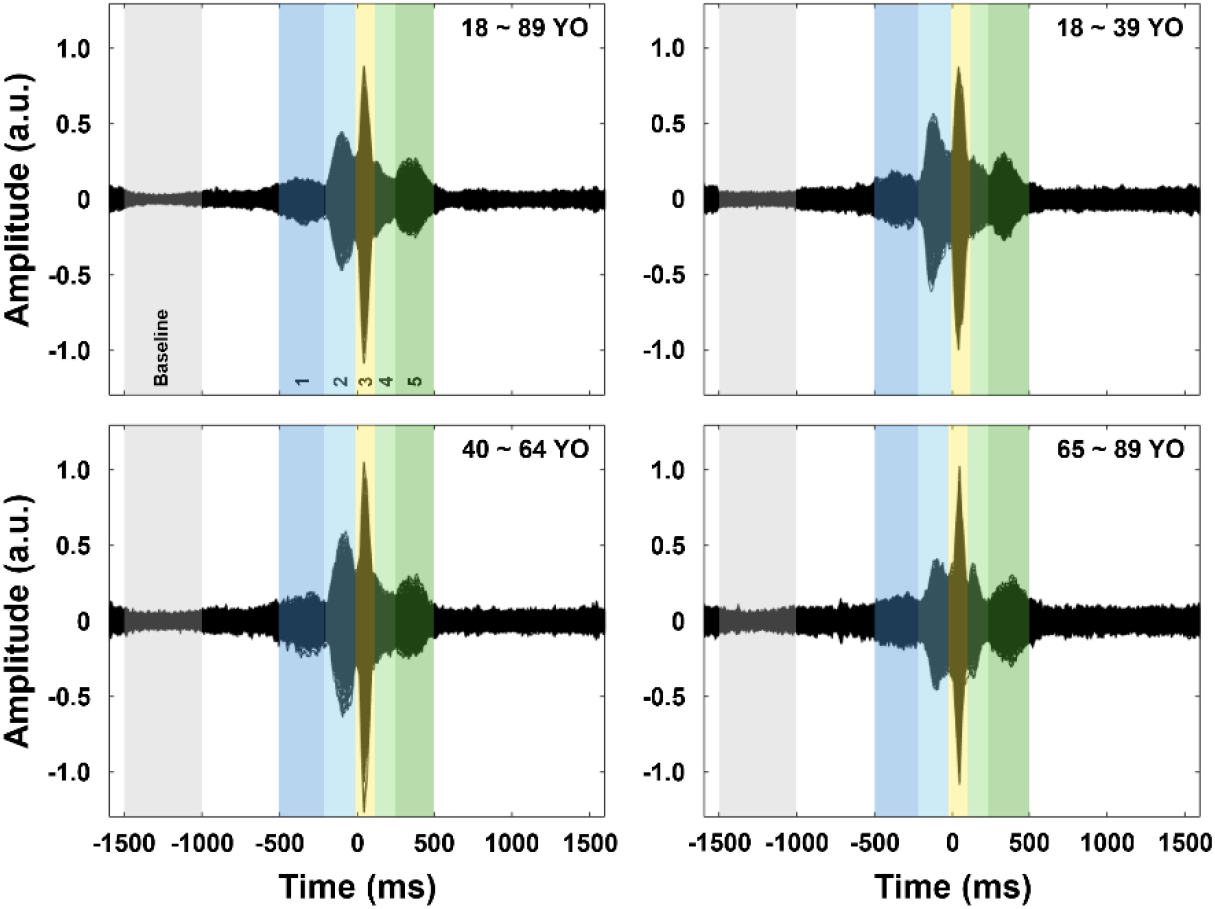
Temporal profiles of group-level cortical neural responses evoked by visual-auditory cued finger tapping. In each plot, black curves represent vertex-wise neural responses time-locked to finger tapping onsets as measured with mean dSPM estimations across trials and across participants, color panels highlight the baseline and the five components as determined by the morphology of the group-level evoked neural responses.

Neural connectivity analysis was performed for ROIs that were revealed from the group-level cortical activation significance testing, which revealed 16 regions with significant activations around the cortical somatosensory and motor areas, 9 regions in the left contralateral hemisphere and 7 regions in the right ipsilateral hemisphere. A ROI was defined as the area covering about 50 vertices that had the strongest activations in a cortical anatomical region as segmented and labeled by the HCP-MMP1 atlas (Glasser et al., 2016). ROIs defined at the group level were projected back to individual cortex surface. Tapping-evoked or RS neural source waveforms (dSPM estimations) for each ROI were extracted from corresponding VACFT trials or RS trials for each individual participant. The extracted ROIs’ source waveforms were further analyzed with the corrected imaginary phase locking value (ciPLV), which is a type of phase synchronization connectivity measure robust to linear transformations of sensor data (Ewald et al., 2012). Computation of pairwise ciPLV was completed by calling the Brainstorm ciPLV connectivity function with Hilbert time-frequency decomposition of time series into six frequency bands including delta (2 ~ 4 Hz), theta (5 ~ 7 Hz), alpha (8 ~ 12 Hz), beta (13 ~ 29 Hz), low-gamma (30 ~ 59 Hz), and high gamma (60 ~ 90 Hz). Pair-wise ciPLV was averaged within the baseline, each of the five components of the tapping-evoked fields, and across the whole trial of RS fields in the six frequency bands. Node strength of connectivity of a ROI that was defined as the sum of ciPLV between the ROI and any ROI among the other 15 ROIs was computed for individual’s tapping-evoked fields and RS fields. Individual participant’s node strength estimations of a ROI were correlated with individual mean target errors during the final adaptation phase with 30° cursor visual feedback rotation. After computation of node strengths, the mean of node strengths was calculated to estimate the SMN global connectivity strength (Wang et al., 2018) and it was correlated with individual mean target errors during final VMA.

### Statistical Analysis

Paired permutation t tests (5,000 permutations) were performed with Brainstorm to test significance of cortical activations underlying tapping-evoked fields. The temporal dimension of the dSPM estimations was reduced to the five components in tapping-evoked fields and the whole trial of RS fields in the aim to control the number of multiple comparisons. Significance level was set at 0.001 (uncorrected for multiple comparisons) to control the rate of type I errors. Correlation between node strength of connectivity of a ROI or SMN global connectivity and mean target error in final VMA was computed with Pearson correlation and partial correlation with age as a covariate. Correlations and partial correlations were deemed as significant if p < 0.001 (uncorrected).

## Results

A total of 96 right-handed participants were sampled from the Cam-CAN repository. Participants selected had MEG recordings during RS and VACFT, brain structural MRI, VMA, and handedness assessment data available. The numbers of females and males were balanced in each of the age groups. Review of MEG data lead to exclusion of 2 participants in the 40 to 59 YO group and 2 participants in the 60 to 89 YO group because of excessive artifacts in their MEG recordings. Data from 92 participants were included in the final analysis. Table 1 listed sample statistics in each of the three age groups predefined.

**Table 1.** Characteristics of the participants included in the final analysis.

| | Age<br>(mean ± SD; years) | Sex<br>(female : male) | Hand dominance<br>(mean ± SD; a.u.) |
| --- | --- | --- | --- |
| 18 to 39 YO | 30 ± 5.9 | 16:16 | 92 ± 12.2 |
| 40 to 59 YO | 51 ± 7.2 | 15:15 | 97 ± 6.3 |
| 60 to 89 YO | 75 ± 5.5 | 15:15 | 96 ± 6.9 |

### Temporal profiles of tapping evoked cortical neural responses

Fig. 1 displayed group-level temporal profiles of tapping evoked cortical neural responses in each of the three age groups and in all participants included in the analysis. Within and across the three age groups, five components were shown in the group-level cortical neural responses. The onset and offset times of each of the five components were very consistent across groups, specifically, with the component 1^st^ roughly arising at −500 ms and ending at −200 ms, the component 2^nd^ from −200 ms to −12 ms, the component 3^rd^ from −12 ms to 112 ms, the component 4^th^ from 112 ms to 244 ms, and the component 5^th^ from 244 ms to 500ms, relative to finger tapping onsets. The component 2^nd^ had a peak onset time at −96 ms. The component 3^rd^ was the most prominent component with a peak onset time at 44 ms.

### Spatial profiles of tapping evoked cortical neural responses

Fig. 2 displayed spatial activation profiles of tapping evoked cortical neural responses. Across the five components from −500 ms to 500 ms relative to finger tapping onsets, significant activations were primarily evoked at pre-central gyrus, post-central gyrus, posterior parietal cortex, temporoparietal junction area, insular lobe, inferotemporal cortex, middle and posterior cingulate areas, and occipital cortex. Specifically, in the component 1^st^, the strongest activations showed at primary visual cortex, temporal lobe, intraparietal sulcus, inferotemporal cortex, temporoparietal junction area, and posterior parietal cortex; in the component 2^nd^, strongest activations existed at the central sulcus area dominantly in the contralateral hemisphere; in the component 3^rd^, prominent activations showed in central sulcus area and posterior parietal cortex bilaterally although the contralateral hemisphere still had a dominance of stronger activations; in the component 4^th^, strongest activations exhibited at insular lobe bilaterally; in the component 5^th^, strongest activations displayed at the post-central gyrus bilaterally with a contralateral dominance. The last plot in Fig. 2 overlayed significant cortical activations across the five components that spanned from −500 ms to 500 ms. Somatosensory and motor areas with strongest activations were selected as ROIs in the following SMN neural connectivity analysis. The spatial localizations of these ROIs were rendered in Fig. 3.

**Fig 2.**
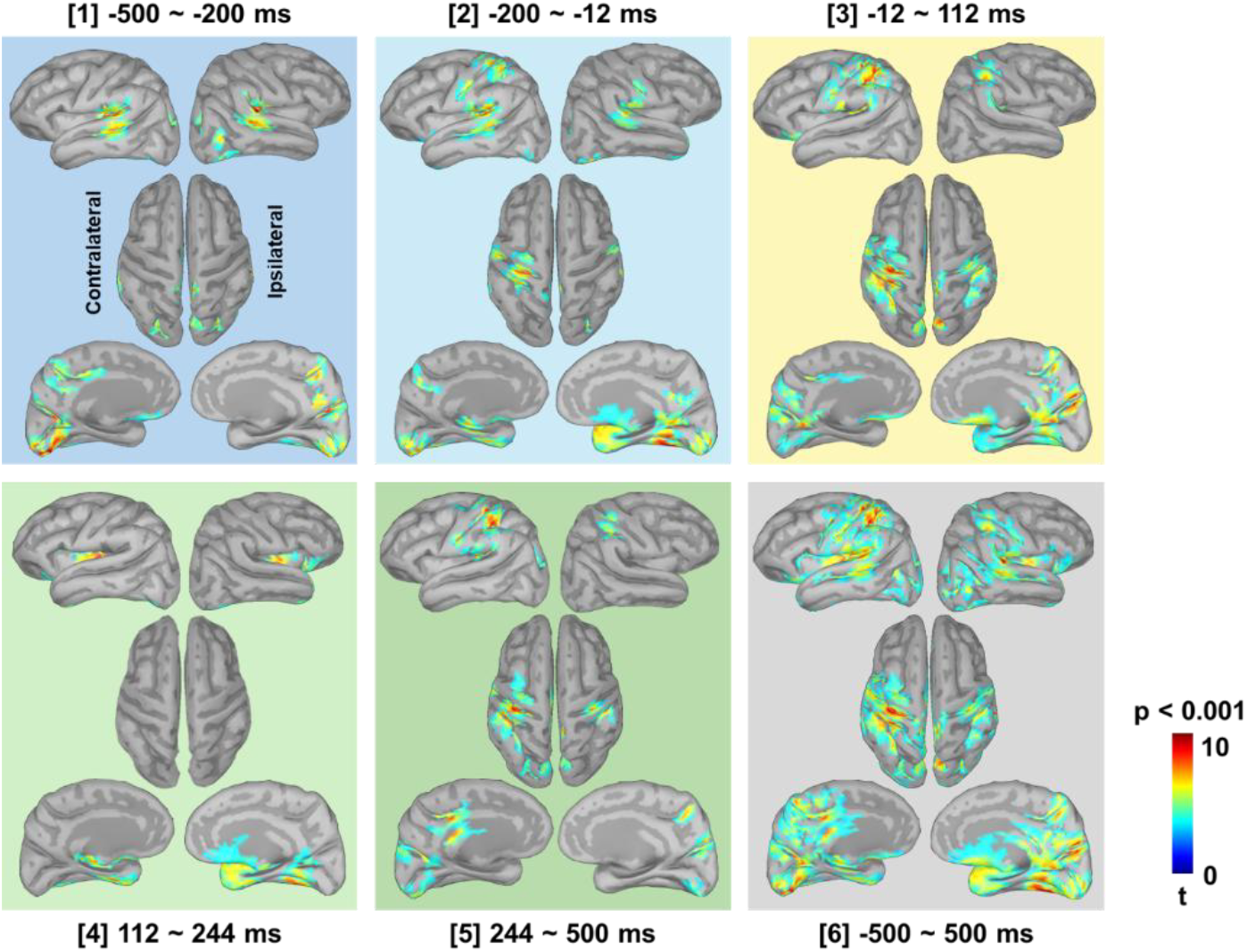
Spatial profiles of group-level cortical neural responses evoked by visual-auditory cued finger tapping. Each plot displayed significant cortical neural activations (p < 0.001 compared to the pre-tapping baseline, uncorrected) corresponding to each of the five components as shown in Fig. 1 and during the time from −500 ms to 500 ms relative to finger tapping onsets.

**Fig 3.**
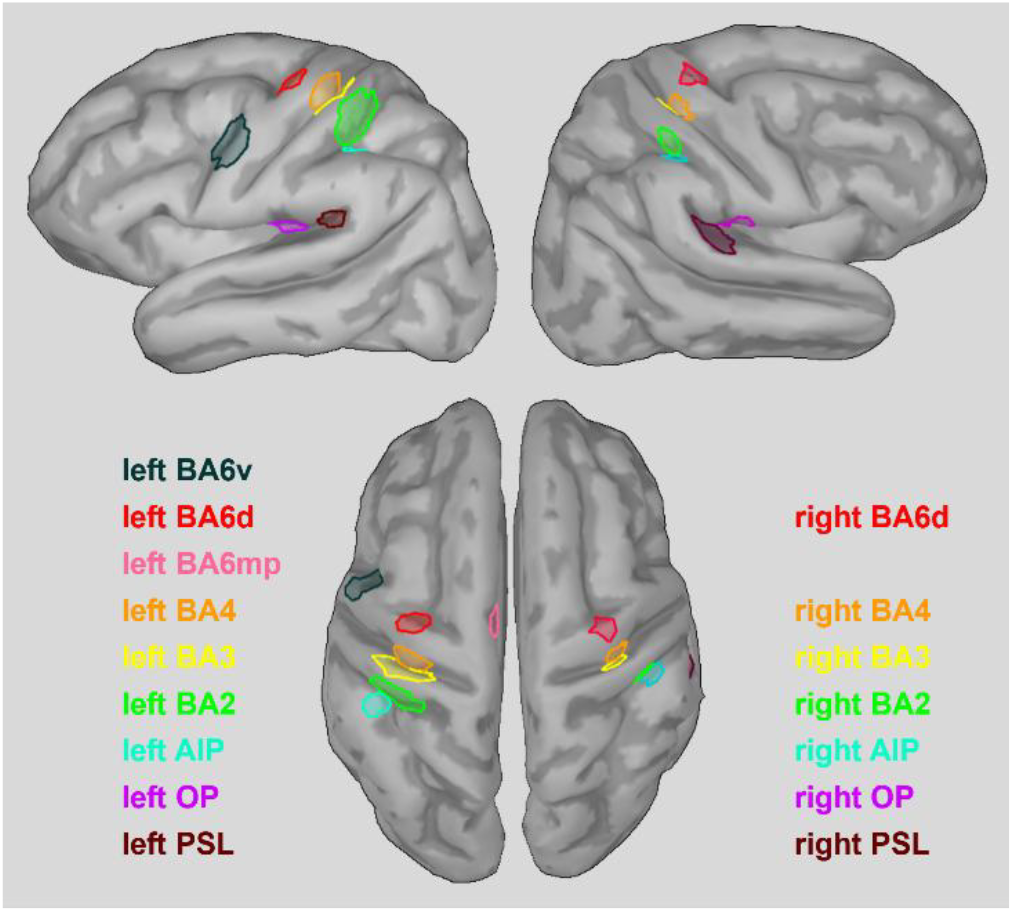
ROIs in the SMN determined by most significant cortical activations during the time from −500 ms to 500 ms relative to finger tapping onsets. BA: Brodmann area; d: dorsal; mp: medial posterior; AIP: anterior inferior parietal; OP: operculum; PSL: perisylvian language.

### SMN global neural connectivity correlations with VMA mean target errors

SMN global neural connectivity strength measured with the mean of ROIs’ node strengths during RS showed no significant correlations or partial correlations with mean target errors in the final 25% of VMA trials (correlation p values > 0.10, partial correlation p values > 0.27, uncorrected). The correlation and partial correlation with age as a covariate between SMN low gamma band global connectivity strength in the VACFT baseline from −1500 ms to −1000 ms relative to finger tapping onsets and VMA mean target errors were approaching significance (r = 0.33, p = 0.0015; partial r = 0.30, p = 0.004), so were the correlation and partial correlation between SMN high gamma band global connectivity strength in the VACFT baseline and VMA mean target errors (r = 0.33, p = 0.0015; partial r = 0.29, p = 0.006). All other correlations and partial correlations between SMN global connectivity strength in VACFT and VMA mean target errors were not significant (correlation p values > 0.01, partial correlation p values > 0.047).

### SMN node specific neural connectivity correlations with VMA mean target errors

Fig. 4 showed significant correlations between node specific neural connectivity in VACFT measured with node strength and mean target errors in the final 25% of VMA trails at the cutoff of p < 0.001. Partial correlation analysis with age as a covariate revealed no significant partial correlations. Specifically, the left BA3 low gamma band node strength in the VACFT baseline was significantly correlated with individual mean target errors (r = 0.35, p = 0.00075; partial r = 0.33, p = 0.0015), so was the left BA6d delta band node strength in the component 1^st^ from −500 ms to −200 ms (r = 0.32, p = 0.0003; partial r = 0.34, p = 0.0011), the left BA6mp high gamma band node strength in the baseline (r = 0.34, p = 0.0009; partial r = 0.32, p = 0.002), and the left BA6mp high gamma band node strength in the component 3^rd^ from −12 to 112 ms (r = 0.35, p = 0.0006; partial r = 0.34, p = 0.001).

**Fig 4.**
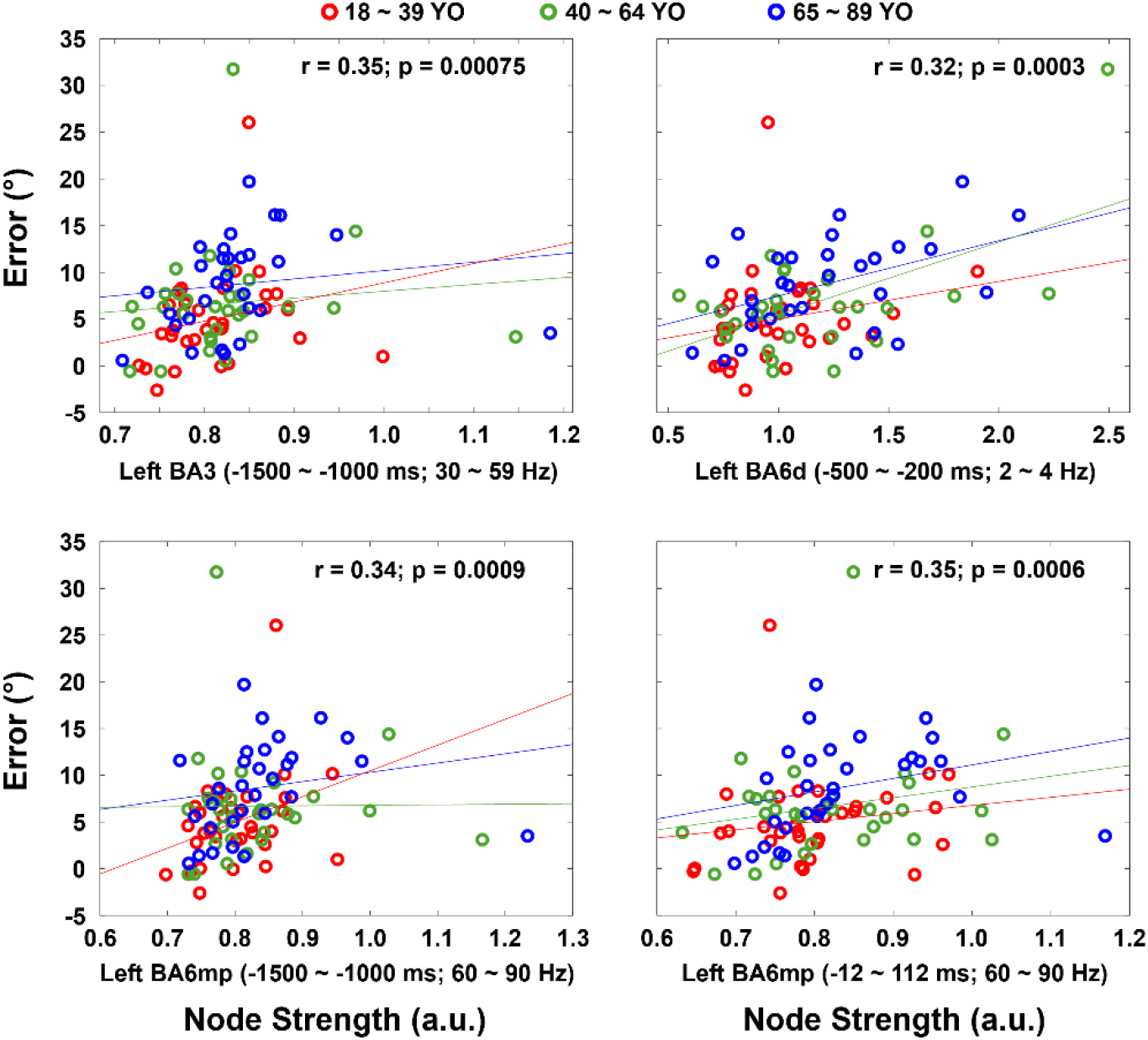
Significant correlations between node strength in tapping-evoked fields and final visuomotor adaptation mean target errors (p < 0.001, uncorrected). The trend line for each group was added only for the purpose of visualization.

Other correlations approaching significance with p values in the range between 0.005 and 0.001 included: the left AIP high gamma band node strength in the baseline (r = 0.30, p = 0.003; partial r = 0.25, p = 0.016), the left BA2 low gamma band node strength in the baseline (r = 0.29, p = 0.004; partial r = 0.25, p = 0.017), the left BA4 low gamma band node strength (r = 0.33, p = 0.0013; partial r = 0.30, p = 0.004) and high gamma band node strength (r = 0.32, p = 0.002; partial r = 0.30, p = 0.004) in the baseline, the left BA6d low gamma band node strength in the baseline (r = 0.32, p = 0.002; partial r = 0.29, p = 0.006), the right BA4 high gamma band node strength in the baseline (r = 0.29, p = 0.0047; partial r = 0.25, p = 0.016), the right AIP high gamma band node strength in the baseline (r = 0.31, p = 0.0025; partial r = 0.27, p = 0.009), the right OP high gamma band node strength in the baseline (r = 0.30, p = 0.004; partial r = 0.25, p = 0.018), and the right BA3 high gamma band node strength in the component 2^nd^ from −200 ms to −12 ms (r = 0.29, p = 0.005; partial r = 0.25, p = 0.016). In contrast to node specific neural connectivity in VACFT, node specific neural connectivity in RS showed no significant correlations with mean target errors in the final VMA (correlation p values > 0.007, partial correlation p values > 0.018).

## Discussion

The current study presents results from analysis of data from the Cam-CAN study. Key nodes in the cortical SMN were initially localized from finger tapping evoked brain magnetic fields as sampled with MEG. Temporal dynamics and spatial source activations underlying finger tapping evoked fields were reported. SMN node specific neural connectivity at the contralateral S1, SMA, and PMd during different stages around finger tapping were significantly correlated with individual final adaptation target errors, with aging effects on these bivariate correlations approaching significance, while SMN neural connectivity during RS were not significantly correlated with individual final adaptation target errors.

### Spatiotemporal profiles of tapping evoked cortical neural responses

Brain sensorimotor control of sensory cued voluntary movements involves a series of integrated dynamical mechanisms including sensory information processing of external sensory cues, sensory-motor transformation and motor planning, prediction of movement-generated sensory consequences, and sensory (especially proprioception) feedback processing (Wolpert & Ghahramani, 2000; Franklin & Wolpert, 2011). The present study characterizations of spatiotemporal profiles of tapping evoked cortical neural responses revealed temporal dynamics and spatial activations associated with sub-processes underlying brain sensorimotor control. The group-level vertex-wise mean dSPM neural response estimations showed five evident components that were highly latency-consistent and amplitude-comparable across 18 ~ 39 YO, 40 ~ 64 YO, and 65 ~ 89 YO. The component 1^st^ primarily was related to visual and auditory processing, since its timing spanning from −500 ms to −200 ms (relative to finger tapping onsets) is compatible with the ranges of simple reaction times in healthy adults (Der & Deary, 2006) and its significant spatial neural activations occurred at primary visual cortex, auditory cortex, intraparietal sulcus, inferotemporal cortex, temporoparietal junction area, and posterior parietal cortex, which locate along the ventral and dorsal visual pathways (Goodale & Milner, 1992; Vaziri-Pashkam & Xu, 2017).

The component 2^nd^ underpinned motor planning related processes, since its timing spanning from −200 ms to −12 ms with a peak response latency at −96 ms, which is consistent with the reported peak latency of motor fields or readiness fields (Cheyne, 2013; Nagamine et al., 1996) and comparable to the reported median onset time (−71 ms) of a change of monkey motor cortex single neuron activity before effector EMG onset (Griffin & Strick, 2020). The component 2^nd^ significant spatial neural activations exhibited dominantly at the contralateral left central sulcus including M1 and premotor cortex that play central roles in planning movements and generating motor commands (Fine & Hayden, 2022). Besides, the temporoparietal junction area also showed significant activations in this component, which is consistent with functions of this area involving assembling multi-modal sensory and motor information (Whitlock, 2017).

The component 3^rd^, the most prominent component, was likely related to somatosensory processing, for that it spanned from −12 ms to 112 ms with a peak response latency at 44 ms. The timing span of this component is consistent with the reported onset time of motor-evoked fields (Cheyne & Weinberg, 1989); the peak response latency at 44 ms is also consistent with the reported peak latency (35.8 ± 9.7 ms) of motor-evoked fields (Onishi et al., 2011). The most pronounced activations showed at the contralateral post-central sulcus area, which is in accordance with previous findings (Onishi et al., 2011) and implies that the component 3^rd^ supported finger tapping related somatosensory processing. The components 4^th^ and 5^th^ may reflect later parts of motor-evoked fields, but they were rarely reported in the literature. Across the five components, key nodes in the SMN were distinguished through their evident activations. The key nodes selected as ROIs to analyze SMN neural connectivity were located within the scope of the SMN mapped with the 7-network parcellation from fMRI in RS (Yeo et al., 2011) as well as the scope of the somato-cognitive action network mapped with fMRI (Gordon et al., 2023).

### Finger tapping evoked SMN neural connectivity correlates with VMA target errors

The contralateral left BA3 low gamma neural connectivity in the tapping baseline (−1500 to −500 ms) was significantly correlated with VMA mean target errors. BA3 is a subarea of primary somatosensory cortex (S1), which has been demonstrated to support formation and consolidation of force-field adaptation and VMA memories in humans in several studies from the Ostry lab. They applied transcranial magnetic stimulation (TMS) to disrupt neural activities at the contralateral S1, M1, and a control site during or after motor adaptation; they reported that disruption of neural activity at S1, but not M1, impaired formation and consolidation of motor adaptation memories (Darainy et al., 2023; Ebrahimi et al., 2024; Kumar et al., 2019). S1 functional connectivity during VMA measured with fMRI was associated with consolidation of VMA memories (Struber et al., 2024). S1 functional connectivity before observation and force field adaptation measured with fMRI predicted individual adaptation performance gains after observation (McGregor & Gribble, 2017). The present finding on BA3 significant correlation between its baseline low gamma neural connectivity and VMA mean target errors further supports that S1 plays critical roles in VMA.

The left BA6mp high gamma neural connectivity in the tapping baseline and the component 3^rd^ (−12 to 112 ms) were significantly correlated with VMA mean target errors. BA6mp, otherwise known as supplementary motor area (SMA), is involved in planning and initiation of voluntary movements, sensorimotor integration, and learning new stimulus-response associations (see review, Nachev et al., 2008). A previous study (Bonzano et al., 2015) reported SMA RS functional connectivity measured with fMRI predicted individual differences in a sequential finger opposition learning task. The present findings on SMA at different tapping components further indicate that SMA may have diverse dynamical functions during VMA, echoed by the finding that SMA activation increased initially and decreased gradually during VMA (Tzvi et al., 2020).

The left BA6d delta band neural connectivity in the component 1^st^ (−500 to −200 ms) was significantly correlated with VMA mean target errors. BA6d covers dorsal premotor cortex (PMd), functions of which include planning sensory triggered voluntary movements, sensorimotor integration, and motor learning (Hoshi & Tanji, 2007; Hardwick et al. 2015). A previous study (Sugiyama et al., 2024) reported that TMS delivered to PMd during motor adaptation impaired meta-learning or decision making to regulate adaptation to maximize rewards, suggesting that PMd may support cognitive-related functions in motor adaptation. The present finding on BA6d in the component 1^st^ aligns with these functions of PMd.

Significant correlations and correlations approaching significance between node specific neural connectivity and VMA mean target errors are frequency specific, mostly shown in the gamma band (low gamma and high gamma) across nodes in the tapping baseline. Gamma band neural activity has been extensively linked to interregional communication and synaptic plasticity (for review, see Fernandez-Ruiz et al., 2023; Buzsáki & Schomburg, 2015), which are fundamental processes underlying learning and memory. Movement-related gamma power increase may reflect intracortical inhibition and be correlated with human motor learning (Zich et al., 2025). Noninvasively modulation of cortical theta and gamma neural coupling promoted motor learning in humans (Akkad et al., 2021). These previous findings and the present findings on SMN gamma neural connectivity support a critical relevance of gamma neural mechanisms in motor learning.

### Resting state SMN neural connectivity not significantly correlate with VMA target errors

Neither SMN global neural connectivity nor node specific neural connectivity in RS was significantly correlated with mean target errors in final VMA. These are different from previously published findings that MEG or EEG RS functional connectivity at the contralateral M1 (Mary et al., 2017; Mottaz et al., 2024; Sugata et al., 2020; Wu et al., 2014) and superior parietal cortex (Manuel et al., 2018) were significantly correlated with individual motor learning performances. Methodological differences between the present study and the published studies may be primary reasons for the discrepancy. First, approaches to estimate functional connectivity are different, previous studies employed a whole brain search approach while the present study focused on the functionally localized SMN. Second, ways to define ROIs are different, previous studies either selected ROIs based on standard anatomical parcellations or based on EEG standard electrode placement montage while the present study defined ROIs based on functional localization via task evoked neural activation significance testing. Third, motor learning tasks are different, previous studies tested non-VMA tasks mostly motor sequence learning while the present study focused on VMA.

### Aging effects

Aging effects were assessed with partial correlations, none of which reached significance at the level of p < 0.001 but some of which approached significance, including baseline gamma connectivity on global SMN connectivity strength and node strength of the left contralateral BA3, BA6mp, and BA4, the left BA6d delta node strength in the first component, and the left BA6mp high gamma node strength in the third component. These correlations are positive, meaning that as people get older, weaker node specific neural connectivity and global SMN connectivity is related to better VMA. This is consistent with the findings that older adults, compared to younger adults, show more widespread brain activations (Seidler et al., 2010) and enhanced motor network connectivity targeting the contralateral M1 (Michely et al., 2018) in motor tasks, reflecting a type of compensatory mechanism associated with aging.

## Conclusions

This study revealed that SMN node specific neural connectivity at different stages around finger tapping were predictive of healthy individual final VMA adaptation performance, especially at the contralateral S1, SMA, and PMd. The results indicate distributed cortical neural correlates with VMA. Further experimental research is needed to probe what specific functional roles these areas may play in the acquisition, consolidation, and retention of motor memories.

## Data and Code Availability Statement

Data in this study were from the Cam-CAN dataset, which is available at the Cam-CAN data access portal: https://camcan-archive.mrc-cbu.cam.ac.uk/dataaccess/.

MEG data analysis was performed with the open-access toolbox Brainstorm, which is available at the toolbox host website: https://neuroimage.usc.edu/brainstorm/.

## Acknowledgements

The author thanks the Cambridge Centre for Ageing and Neuroscience (CamCAN) for Data collection and sharing and granting use of the data that were analyzed in this study.

CamCAN funding was provided by the UK Biotechnology and Biological Sciences Research Council (grant number BB/H008217/1), together with support from the UK Medical Research Council and University of Cambridge, UK.

